# Long read sequencing of retinal RNA improves killifish transcriptome annotation

**DOI:** 10.64898/2026.09.02.748420

**Authors:** Sohini Rebba, Lianri van Schalkwyk, Aleksandra M Krzywańska, Ryan B. MacDonald, Brian S. Clark, Philip A. Ruzycki

**Author notes:** Corresponding authors: Brian S. Clark, Philip A. Ruzycki.

## Abstract

**Purpose:** The African Turquoise Killifish has recently emerged as a powerful model for aging and age-related disease research studies. However, molecular based investigations have been limited by preliminary genome and transcriptome builds with incomplete reference genome sequence, fragmented chromosome assembly, and missing gene annotations. These issues make primary (alignment and quantification) and secondary (Gene Ontology, Gene Set Enrichment Analysis, cross-species comparisons) analyses difficult to reliably implement and interpret. This study seeks to generate a complete retinal reference transcriptome to facilitate future killifish transcriptomic, epigenetic, and proteomic studies of the visual system.

**Methods:** We generated an enhanced retina transcriptome using long-read PacBio RNAseq data that was processed using a robust computational pipeline to merge reads, classify genes, and annotate with nearest orthologous gene names from other species. This new annotation was compared to available references and validated using bulk and single cell RNAseq datasets.

**Results:** Comparison of the widely used Nfu_20140520 and the newly released NfurGRZ- RIMD1 genome builds identified NfurGRZ-RIMD1 to be more contiguous and complete. However, we identified limitations with both transcriptomes, including the lack of annotation of certain retina specific genes and many uninformative gene names. Using long-read PacBio sequencing of RNA collected from young and old Killifish retinas, we annotated a deep retinal transcriptome onto the NfurGRZ-RIMD1 reference genome. This analysis identified thousands of previously unannotated transcripts from retinas of young and old killifish. By matching each translated protein sequence to its nearest ortholog, we increased the number and proportion of genes with meaningful gene names. Mapping of bulk and single-cell RNAseq data showed substantial increase in mapping rate and identified hundreds of genes and transcripts with age-dependent expression dynamics.

**Conclusions:** Assembly of an enhanced retinal transcriptome for the killifish improved both primary and secondary analyses of bulk and single cell RNAseq data. Improvements will benefit future studies investigating the mechanisms of aging in the killifish and to best utilize this powerful model to understand human disease.

## Introduction

Age is a common risk factor for many diseases of the eye including age-related macular degeneration, glaucoma, cataract, and diabetic retinopathy [1–3]. The vision research community has found numerous effective models for rare inherited ocular diseases [4,5] and developed targeted experimental conditions to mimic specific phenotypes of common, complex age-associated diseases [5–7]. However, experimentally modeling organismal aging in traditional model organisms is challenging due to prolonged aging timescales, high costs, and other practical limitations. Therefore studying organisms that have shorter lifespans and display age-related pathology and dysfunction is essential to test causative cellular mechanisms of aging diseases, guiding the development of novel therapies to slow or reverse these devastating conditions.

The African Turquoise Killifish (*Nothobranchius furzeri*) has the shortest natural lifespan amongst vertebrates that propagate within the laboratory (4-6 months) and displays numerous hallmarks of aging across tissues [8–10]. As a model, the killifish can be maintained within a laboratory setting with only select modifications from traditional zebrafish husbandry procedures [11]. The killifish has a sequenced genome and is amenable to genetic manipulation by tools such as CRISPR [12–16], making it a powerful new laboratory model for aging and age-related disease research [17].

The killifish retina displays many hallmarks of aging including excess DNA damage, senescence markers, reactive gliosis, and decreased visual acuity [18]. Recently, we published the first bulk and scRNAseq analyses over the killifish retina lifespan [19]. While the data identified genes that change expression over time by both modalities, an exhaustive or cross-species integrative analysis was hampered by relatively low mapping rate of both modalities to the reference transcriptome and an inability to utilize unbiased tools like gene ontology (GO), Gene Set Enrichment (GSEA), or others to better understand the global functional relevance of modified gene sets. Additionally, as aging has been linked to epigenetic changes over time, we reasoned that future efforts to integrate transcriptomic and epigenomic datasets would be restricted by both the genome build and transcriptome reference quality. Therefore, efforts to improve these essential tools is warranted to maximize utility of the killifish to study genetic mechanisms of age-related disease research.

Published transcriptomic studies have largely used the killifish genome and transcriptome that were released in 2014 (Nfu_20140520) [14,19–24]. Our published work on the killifish retina utilized this reference genome but was enhanced by integration of a custom transcriptome build based on RNA from the telencephalon [25]. While there is no specific cellular overlap between the retina and telencephalon, the neural optimized transcriptome increased mapping rates of retina bulk and single-cell RNA data, suggesting some genes or isoforms were not present in standard reference files [19]. More recently a new reference (NfurGRZ-RIMD1) was released, though uptake of this new tool has been limited to date [26]. In this work, we sought to compare the two reference genomes available for the killifish and annotate a new, complete killifish retinal transcriptome to enhance future research assessing transcript expression within this important new model.

## Methods

### Killifish husbandry

GRZ strain turquoise killifish (*Nothobranchius furzeri*) were housed at University College London under a 14:10 light:dark cycle on PPLs PP2133797. Killifish were hatched once they reached the golden eye stage by placing the embryos in an aerated 1:1 mixture of 4°C hatching solution (1g/L humic acid in autoclaved MilliQ water):room temperature system water. Adult fish (4 weeks post hatching) were fed dry food and bloodworm. For all experiments, “young” fish were considered 6 to 8-week-old young adults, and “old” fish were considered 22 to 24-week-old adults at the end of their lifespan.

### Sample collection and processing

Retinas were extracted and placed into 1ml Trizol before being frozen and stored at - 80C. RNA was isolated based on published protocols [27]. Briefly, retinas were homogenized with a motorized pestle before addition of 200ul chloroform. Samples were then vortexed and centrifuged for 5min at max speed. The aqueous layer (∼700ul) was transferred to a new tube. Equal volume 70% Ethanol in DEPC-treated H2O was then added before the entire sample was loaded onto a Zymo Research RNA clean and concentrator column. RNA was subjected to in-column DNAse treatment, washed and eluted per manufacturer’s protocol. RNA yield was approximately 450ng/ul for each sample and stored at -80C prior to library preparation.

### Library preparation: PacBio long-read RNA sequencing

Long-read RNA sequencing was performed using the PacBio Kinnex Full-Length RNA sequencing protocol at the Genome Technology Access Center (GTAC), McDonnell Genome Institute, Washington University in St. Louis. Two total retinal RNA samples from young (8-week-old) and old (22-week-old) African turquoise killifish were submitted for sequencing. The Kinnex protocol uses 300 ng of total RNA as input and captures polyadenylated transcripts, thereby excluding ribosomal RNA during library preparation. Individual RNA samples from 8- and 22-week-old retinas were converted into individually barcoded cDNA libraries and subsequently pooled into a single Kinnex library for sequencing on one PacBio Revio SMRT Cell. The Kinnex workflow concatenates multiple transcript molecules into a single HiFi sequencing read prior to sequencing. Following sequencing, PacBio Read Segmentation was used to separate HiFi reads into individual full length transcript reads (segmented reads, S-reads). The resulting segmented BAM files were used as the input for downstream transcriptome building.

### Demultiplexing of Kinnex reads

Segmented reads generated from the Kinnex workflow were demultiplexed using PacBio lima (v2.13.0) with –isoseq and --peek-guess to identify and remove sample specific barcodes and adapter sequences to generate demultiplexed and trimmed reads.

### Iso-Seq processing

Lima processed reads were analyzed using the Iso-Seq refine tool to identify FLNC (Full-Length Non-Chimeric) transcript reads, containing both 5’ and 3’ cDNA primers with the polyA tail to trim. FLNC reads were clustered using Iso-Seq cluster2 to generate high quality consensus transcript sequences. Consensus sequences were aligned to the NfurGRZ-RIMD1 genome build using pbmm2 (--isoseq present). Redundant transcript isoforms were collapsed using Iso-Seq collapse to generate a nonreductant transcriptome.

### SQANTI3- Quantity control and transcript classification

The collapsed transcriptome was analyzed using SQANTI3 [28] to classify transcript isoforms relative to NCBI reference annotation. SQANTI3 (sqanti3_qc.py) integrates splice junction support, transcript structural classification and additional quality assessment to identify known and novel isoforms. Published killifish bulk RNAseq data [22] was also incorporated to improve transcript quality by filtering transcripts based on short read coverage support. Finally, transcripts were filtered based on the number of exons to remove mono-exonic transcripts (sqanti3_filter.py).

### Functional annotation

The SQANTI3 filtered transcriptome was analysed using TransDecoder [29] to predict open reading frames (ORFs) for all transcript isoforms. Predicted ORFs were annotated using EggNOG-mapper [30], which assigned orthologous gene names and functional annotation based on sequence homology match to *Actinopterygii* (Ray-finned fish) as the closest match in the EggNOG database. The resulting annotations were manually curated to resolve instances where identical gene names were assigned to distinct genomic loci by appending locus-specific identifiers thereby maintaining unique gene names while preserving compatibility with previous killifish transcriptome annotations (e.g. H3F3A_1, H3F3A_2). Gene name annotations from EggNOG are reported exactly as returned by the analysis and as such all gene names do not conform to traditional fish nomenclature (lowercase and italicized).

### Bulk RNA-seq analysis

Previously published bulk RNA-seq datasets were re-mapped to the Nfu_20140520 and NfurGRZ-RIMD1 genome builds to compare performance of different transcriptome annotations. Identical quantification and filtering parameters were applied across the three transcriptomes to compare mapping performance, gene and transcript detection, differential expressions and other downstream analysis. For gene-level analysis, short-read RNA-seq reads were aligned using STAR (v2.7.0d) [31]. Gene level counts were generated using featureCounts (v2.0.0) [32]. Differential expression analysis was performed using DEseq2 (v1.44.0) [33]. For transcript level analysis, transcript abundance was quantified using Salmon (v1.10.3) [34]. Salmon outputs were imported into R using tximport (v1.32.0) [35] and downstream analysis was performed using DESeq2 (v1.44.0).

### Single-cell RNA seq analysis

Previously published single cell RNA-seq data were re-mapped using Cellranger (v9.0.1) with references generated from Nfu_20140520, NfurGRZ-RIMD1 and PacBio transcriptome annotation. Identical processing and filtering parameters were applied across three references to compare mapping performance, gene detection and downstream single-cell analysis using Seurat (v5.4.0) [36].

### Hybridization Chain Reaction

In situ hybridization chain reaction (HCR) was carried out on slides following previously published protocols [37] with minor modifications. Probes were designed against PB.6621 and PB.6619 using a custom python script from the Seuntjens lab [38,39] (Github: https://github.com/SeuntjensLab/Easy_HCR) and obtained from ThermoFisher. The slides were pre-hybridized in a pre-warmed probe hybridization buffer (Molecular Instruments, www.molecularinstruments.com) for 30 minutes at 37°C and probe solutions were prepared to a final concentration of 24pmol/mL in the probe hybridization buffer. Probe mix was added to the slides and allowed to incubate for 3 days in a humidity chamber at 37°C. To remove excess probe solution, the slides were washed at 37°C in probe wash buffer (Molecular Instruments) 3 x 10 minutes and then in 5x SSCT at room temperature for 2 x 5 minutes. Amplifiers were prepared by individually heating 6pmol of each of the H1 and H2 hairpins from HCR B2-647-conjugated and B3-546-conjugated amplifier sets (Molecular instruments) at 95°C for 90 seconds. The hairpins were then snap cooled in the dark before being added to the amplification buffer (Molecular Instruments). Slides were pre-amplified in an amplification buffer for 30 minutes at room temperature and the amplifier solution was added to the slides and left overnight in the dark in a humidity chamber. The slides were washed at room temperature in 5x SSCT 4 x 15 minutes, stained with DAPI and mounted using FluoromountG (Thermo Fisher Scientific). Probe sequences for PB.6621 (*rh2.2*) and PB.6619 (*rh2*) are provided (**Supplementary_Methods.xlsx**).

### HCR Imaging

Imaging was performed using a Zeiss LSM 900 inverted confocal microscope with a 40X water-immersion objective. For super resolution imaging of *in situ* HCR, imaging was performed using the Zeiss 4Y-Airscan setting. To capture the full retinal thickness, z-stacks were acquired at 0.3*μ*m intervals. Imaging was performed on the central retina, defined as the midpoint between the two opposing ciliary marginal zones within sections.

### Data and code availability

The PacBio Kinnex long-read RNA sequencing data generated in this study is available in NCBI sequence Read Archive (SRA) under BioProject accession **PRJNA1511413** (SRA run accession: **SRR40124706**). The improved killifish transcriptome (**GSE344270_Retina_Rebba.gtf.gz**) is available in the NCBI Gene Expression Omnibus (GEO) under accession **GSE344270**. The computational workflow and the processing parameters used to generate the transcriptome is available at our Github repository: https://github.com/p-ruzycki/killifish-pacbio-transcriptome. Genome assemblies and reference transcriptome annotations used were accessed via the following urls: Nfu_20140520:https://nfingb.leibniz-fli.de/; GCF_043380555.1_NfurGRZ- RIMD1:https://www.ncbi.nlm.nih.gov/datasets/genome/GCF_043380555.1/.

## Results

### Comparison of existing genome builds

Multiple killifish reference genomes are available but no thorough comparison has yet been reported [24,26]. Nfu_20140520 is still commonly used for transcriptome analyses across tissues of the killifish [14,19–24]. In 2025, a new build (NfurGRZ-RIMD1) was released by the Squalomix Consortium [26], though to date there has not yet been a publication describing the build or any comparison to the Nfu_20140520 genome. Few studies have published results utilizing the new NfurGRZ-RIMD1 build, though some have switched [40]. Therefore, we first sought to determine which is the more complete reference for transcriptomic and epigenetic studies. We utilized BlobToolKit [41] to quantify and visualize metrics of chromosome length, genome size, nucleotide composition, and other important metrics (**Figure 1A-B**). The NfurGRZ-RIMD1 genome includes nearly 20% more genetic material (1.48G vs 1.24G), chromosomes are more complete (higher N50, N90, and auN), and the NfurGRZ-RIMD1 contains no unannotated bases (N bases 0% vs 31%). Blobtoolkit also assigned the NfurGRZ-RIMD1 assembly a substantially higher overall assembly score (0.66 vs 0.24). Collectively, these results demonstrate that the NfurGRZ-RIMD1 assembly provides a more complete and contiguous genomic reference for downstream transcriptome reconstruction and analysis.

**Figure 1:**
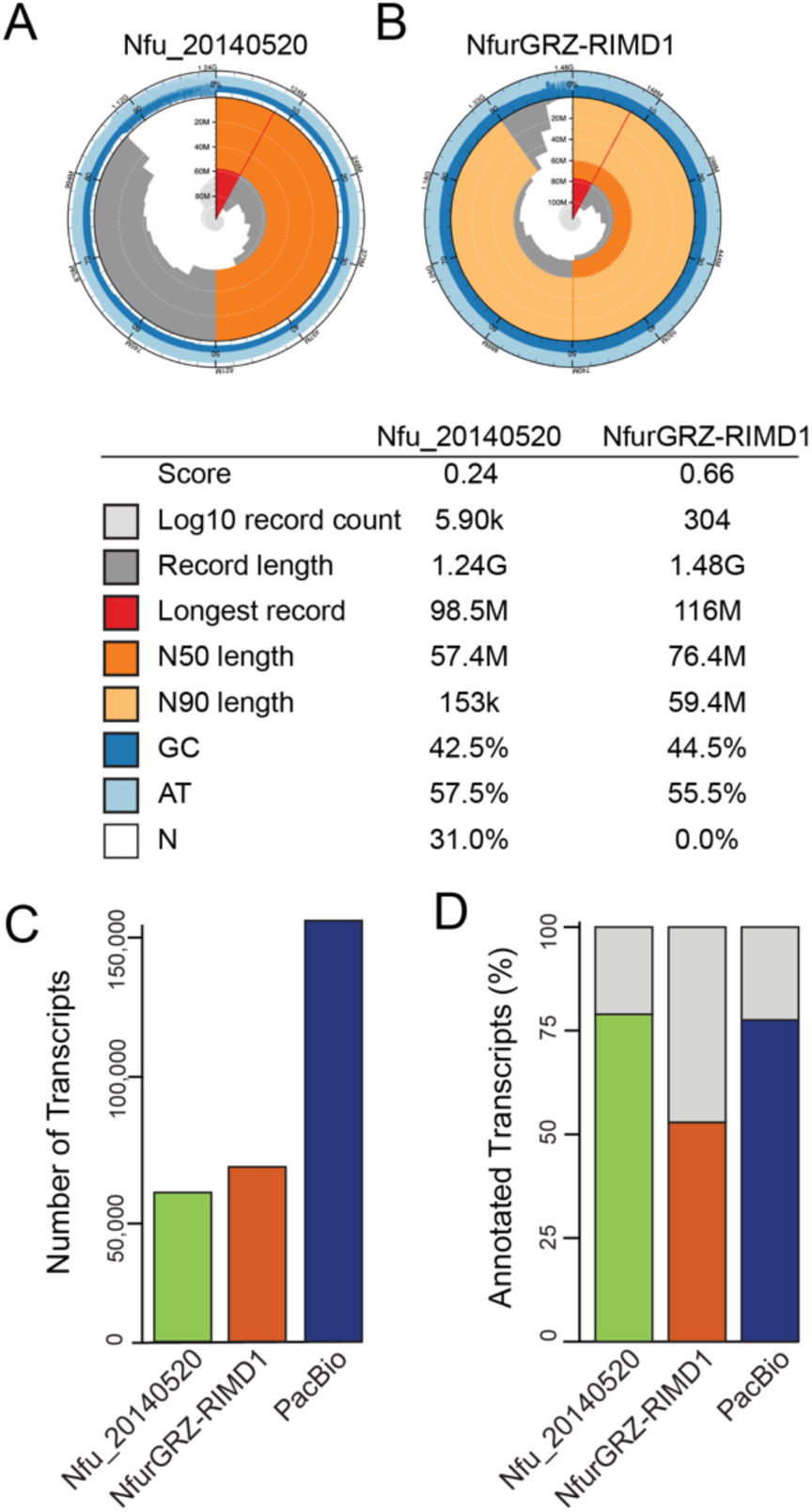
Comparison of killifish genome assemblies and transcriptome quality. (**A-B**) BlobToolKit snail plots comparing the genome assembly statistics of NfurGRZ-RIMD1 and Nfu_20140520 killifish reference genomes. Assembly metrics include total genome size (outer circle), chromosome/scaffold length distribution (grey circle), N50 (dark orange), N90 (light orange), auN (dark grey), longest sequence (red), nucleotide composition (blues) and proportion of unresolved bases (white) and overall assembly score. (**C**) Comparison of total number of transcripts identified in the Nfu_20140520, NfurGRZ-RIMD1 and PacBio transcriptome annotations. (**D**) Comparison of transcriptome annotation across builds, showing the percentage of transcripts assigned biologically meaningful gene names (colored portion of the bars) and transcripts lacking gene name or assigned placeholder names (grey portion of the bars).

However, analysis of the reference transcriptomes revealed a notable limitation of the NfurGRZ-RIMD1 annotation (**Figure 1C-D**). Although the NfurGRZ-RIMD1 reference had substantially more annotated genes than the Nfu_20140520 annotation (38,438 vs 26,739), fewer than 53% of genes were assigned biologically meaningful gene names compared to nearly 70% in the Nfu_20140520 annotation. This reduction was largely driven by the high proportion of the ‘LOC###’ placeholder gene names in the NfurGRZ- RIMD1 reference (47%) whereas place holder or accession names accounted for only 11% of genes in the Nfu_20140520 annotation. A similar trend was seen at the transcript level; while both transcriptomes have roughly equivalent numbers of annotated transcripts (62,661 vs. 59,752), only 53% of NfurGRZ-RIMD1 transcripts were assigned informative gene names compared with 78% of transcripts in the Nfu_20140520 annotation.

### Pipeline and transcriptome annotation stats

As the NfurGRZ-RIMD1 genome build is an improvement but lacks only in the quality of the reference transcriptome, we hypothesized that refinement and re-annotation by integrated analysis of PacBio based Iso-Seq long read RNAseq data and other bioinformatic tools would benefit future studies (**Figure 2**). Total RNA isolated from young and aged killifish retina was used to generate independent PacBio Kinnex full length RNA libraries. The two barcoded libraries were pooled and sequenced on a single PacBio Revio SMART Cell to generate 5,666,376 HiFi reads. Raw data was processed by Kinnex read Segmentation to produce 44,012,991 segmented full length reads (S-reads) with a mean length of 1,893bp. S-reads were processed using the Iso-seq pipeline to generate high-quality, full-length transcript isoforms, resulting in 734,231 retained transcripts. Further refinement by SQANTI3 [28] with integration of 37 published retina bulk RNAseq datasets [22] across the killifish lifespan to validate splice junctions and transcript support resulted in 21,014 genes and 165,015 high confidence transcripts, more than 53% of which were identified as ‘Novel Isoforms’ (**Figure 1C**). This final reference transcriptome was annotated using TransDecoder [29] to predict ORFs and EggNOG [30] to assign orthologs and functional annotations. Functional annotation assigned orthologous gene names to 76.6% of transcript isoforms and the proportion of genes with biologically meaningful gene names was increased from 52.9% to 71.9% while the ‘LOC###’ placeholder gene identifiers were reduced from 47.1% to less than 1% compared to the NCBI reference (**Figure 1D**). Together, this new resource offers a more functionally informative and biologically interpretable reference transcriptome for the killifish retina.

**Figure 2:**
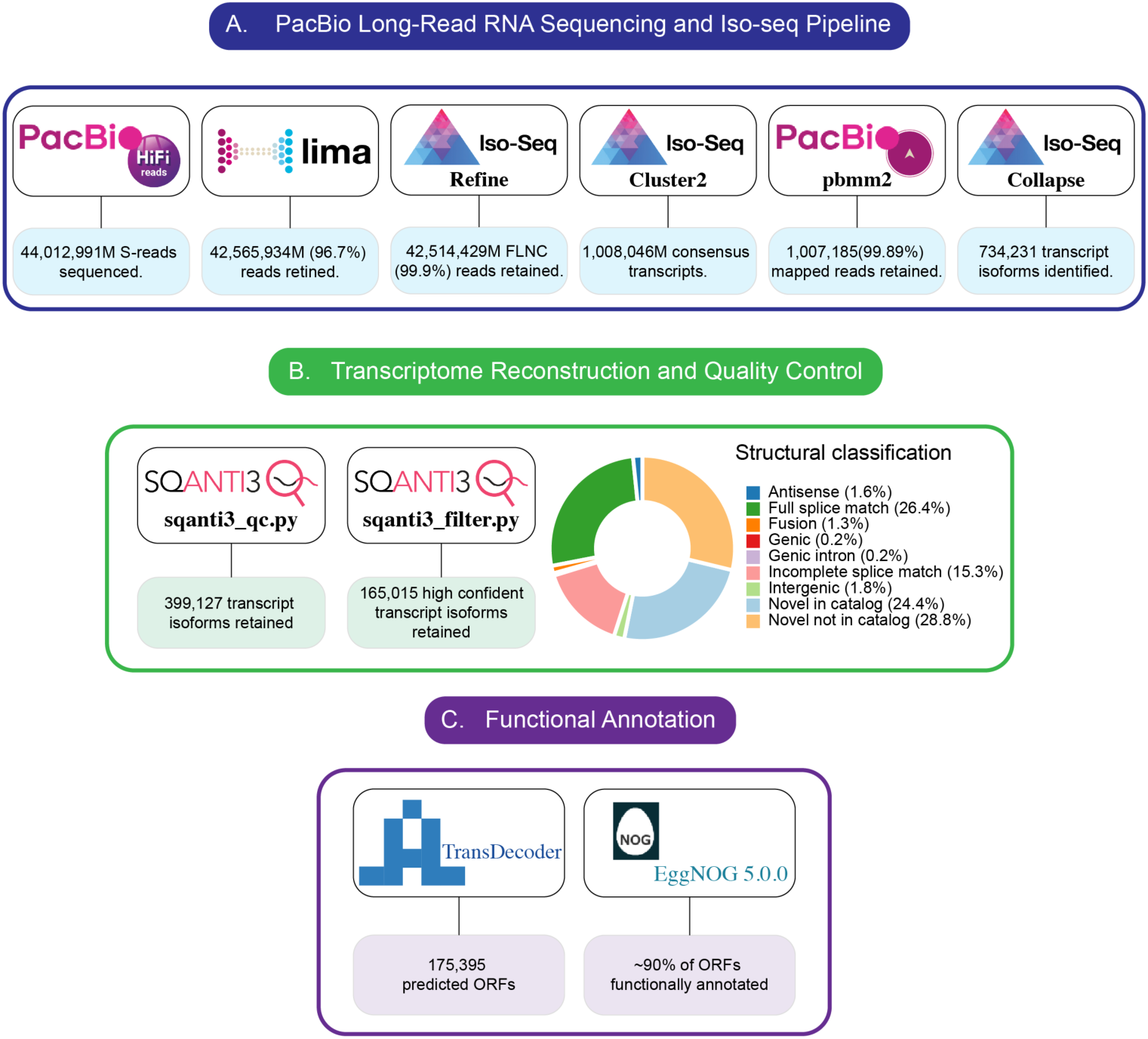
Computational Workflow for PacBio Long-Read Transcriptome Reconstruction and Annotation. (**A**) PacBio Kinnex reads from young and aged killifish retinas were processed through Iso-Seq pipeline, followed by genome alignment using pbmm2 to NfurGRZ-RIMD1 build and isoform collapsing. (**B**) Transcript models were classified and filtered using SQANTI3 incorporating 37 published retinal RNA-seq datasets across the killifish lifespan. (**C**) High confidence transcripts were analysed using TransDecoder for ORF prediction and EggNOG mapper for functional annotation. Numbers indicate reads or transcripts retained at each step of the pipeline.

### Bulk RNAseq remapping and analysis

To validate the utility of the new transcriptome build we reanalyzed previously published bulk RNA seq datasets profiling 14 tissues across the killifish lifespan [22]. Although the transcriptome was generated from retinal long read RNA sequencing it improved mapping rates across the majority of tissues. The largest increases were observed in neural tissues including retina, brain, and spinal cord and mapping rate was only notably worse within liver, fat, and gut (**Figure 3A**).

**Figure 3:**
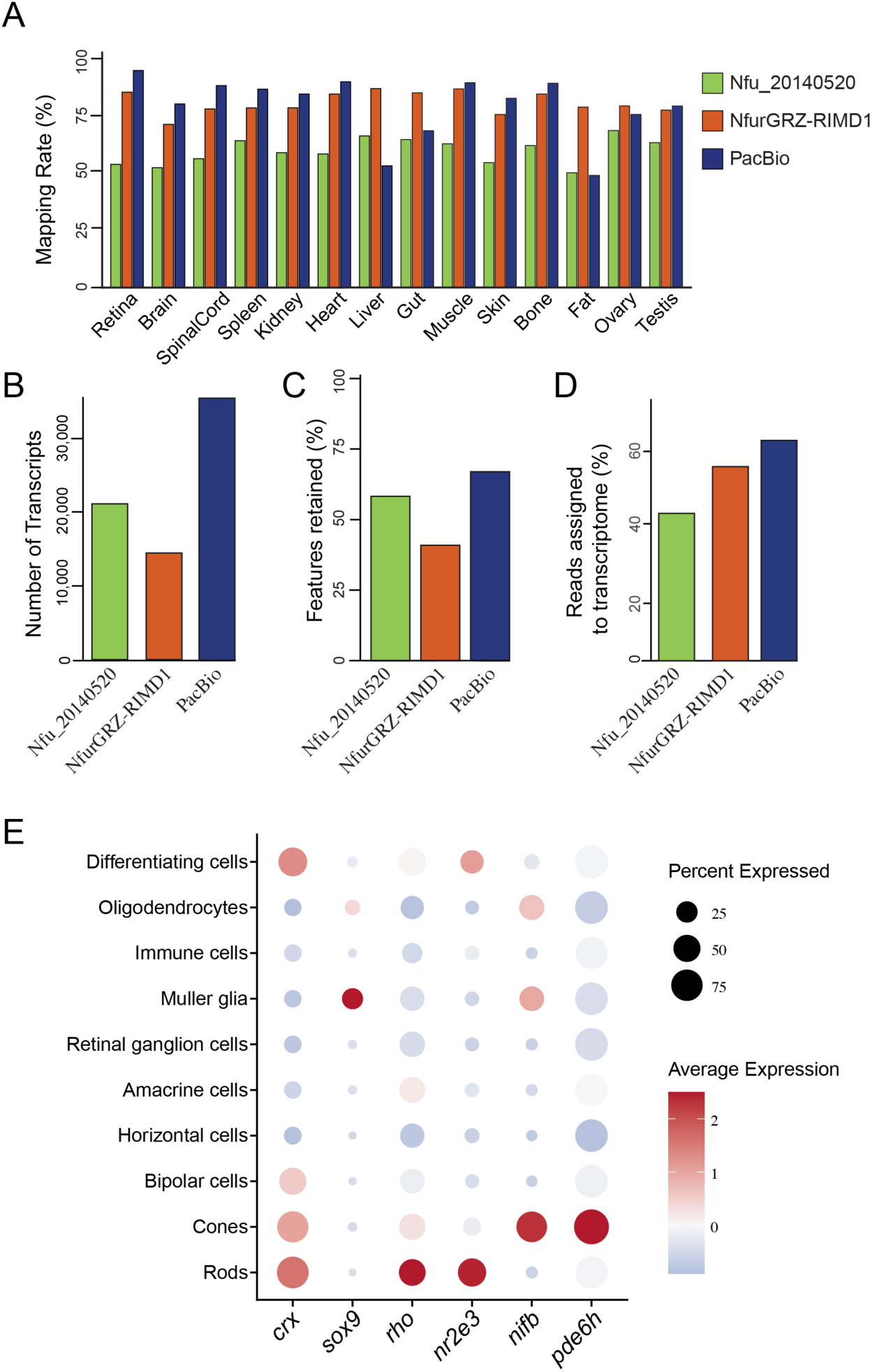
Evaluating Improved Performance of the PacBio Transcriptome in Bulk and Single-Cell RNA-seq Analysis. (**A**) Salmon mapping rates (mean of 1 mapped young and 1 mapped old sample) of published bulk RNA-seq datasets across 14 killifish tissues using the Nfu_20140520, NfurGRZ-RIMD1 and PacBio transcriptomes. (**B**) Number of transcript isoforms retained across retinal ages using a TPM >3 threshold. (**C**) Proportion of genes retained after standard expression filtering of CPM >3 across transcriptome annotations. (**D**) Comparison of transcriptome mapping rates of published retinal scRNA-seq datasets (6-,12-,18-week-old killifish). (**E**) Cell-type specific expression of retinal and photoreceptor genes annotated by the PacBio transcriptome and visualised in remapped scRNA-seq data.

Further analysis focused only on the performance of the PacBio transcriptome for retina datasets. Compared to both Nfu_20140520 and NfurGRZ-RIMD1 annotations, the PacBio transcriptome retained approximately two-fold more expressed transcript isoforms across all ages (TPM >3) showing improved transcript representation within the retina (**Figure 3B**). At gene level, the PacBio transcriptome showed a higher proportion of genes retained after standard filtering (CPM >3), with 66.1% compared to 40.4% for NfurGRZ-RIMD1 and 57.5% for Nfu_20140520 builds (**Figure 3C**).

To further evaluate the performance of the PacBio transcriptome in single cell RNAseq analysis, we re-mapped previously published data profiling 6-, 12- and 18-week-old killifish retinas [19]. Across all samples, the proportion of reads assigned to the transcriptome (transcriptome mapping) improved 6-8% compared to NfurGRZ-RIMD1 and 10-12% compared to Nfu_20140520 builds (**Figure 3D**). Further analysis revealed the PacBio transcriptome retained all previously annotated and even added 24-26% more cells per sample, increasing power for downstream analyses (**Supplementary Figure 2A**).

In our previous analysis of retina RNA-seq data we noticed that some key photoreceptor, retina development, and disease-related genes were not annotated, presenting issues in identification of cell types by scRNA-seq and interpretation of the phenotype of photoreceptor function in aging. Notable examples included a lack of clearly annotated genes in one or both available transcriptomes for *crx*, *nr2e3*, *nfib*, *pde6h*, *sox9*, and even *rho*. All of these are now annotated within our new transcriptome build and analysis with published scRNAseq data clearly shows the expected pattern of strong isolated cell type specific expression (**Figure 3E**).

### Differential expression analysis and validation

We chose to further investigate age-related transcriptomic changes at the gene level. We compared old vs young (21- vs 7-week-old) killifish retina samples from an independent study [22] and identified 1,683 differentially expressed genes (**Figure 4A - B**). Notably, 1,349 of 1,683 (80.2%) have an assigned gene name orthologous to human and other model species (**Supplementary table 1**), comparable or better than similar analyses run with the previous transcriptome and genome references (**Table 1**). Genes identified and *in vivo-*validated for age-dependent transcriptional dynamics in our previous work also displayed statistically significant gene expression changes within the independent 21- vs 7-week-old sample cohort, including *optn*, *nfkb2*, and *pcdh15*. Other genes identified in our studies including *tgfb3*, *apoe*, *htra1*, and *bdnf* did not pass statistical thresholds in this independent dataset and analysis; however, all genes still showed a congruent trend of gained or lost expression with age across studies. We noted that the PacBio transcriptome identified less ‘statistically significant’ genes changing with age compared to the other builds. Unfortunately a direct comparison between the separate lists is not possible. Such discrepancies would be expected with the dramatic differences in gene number, mapping rate, and consistency of transcripts in the reference (**Figure 3B-D**). Lastly, gene ontology analysis of all up- and down-regulated genes identified numerous biological processes, molecular function and cellular component gene sets with significant enrichment. Notably, upregulated sets of genes included genes involved in inflammation, angiogenesis, and extracellular matrix remodelling (**Supplementary Figure 1A**), while downregulated genes showed evidence of cellular metabolism and mitochondrial function (**Supplementary Figure 1B**).

**Figure 4:**
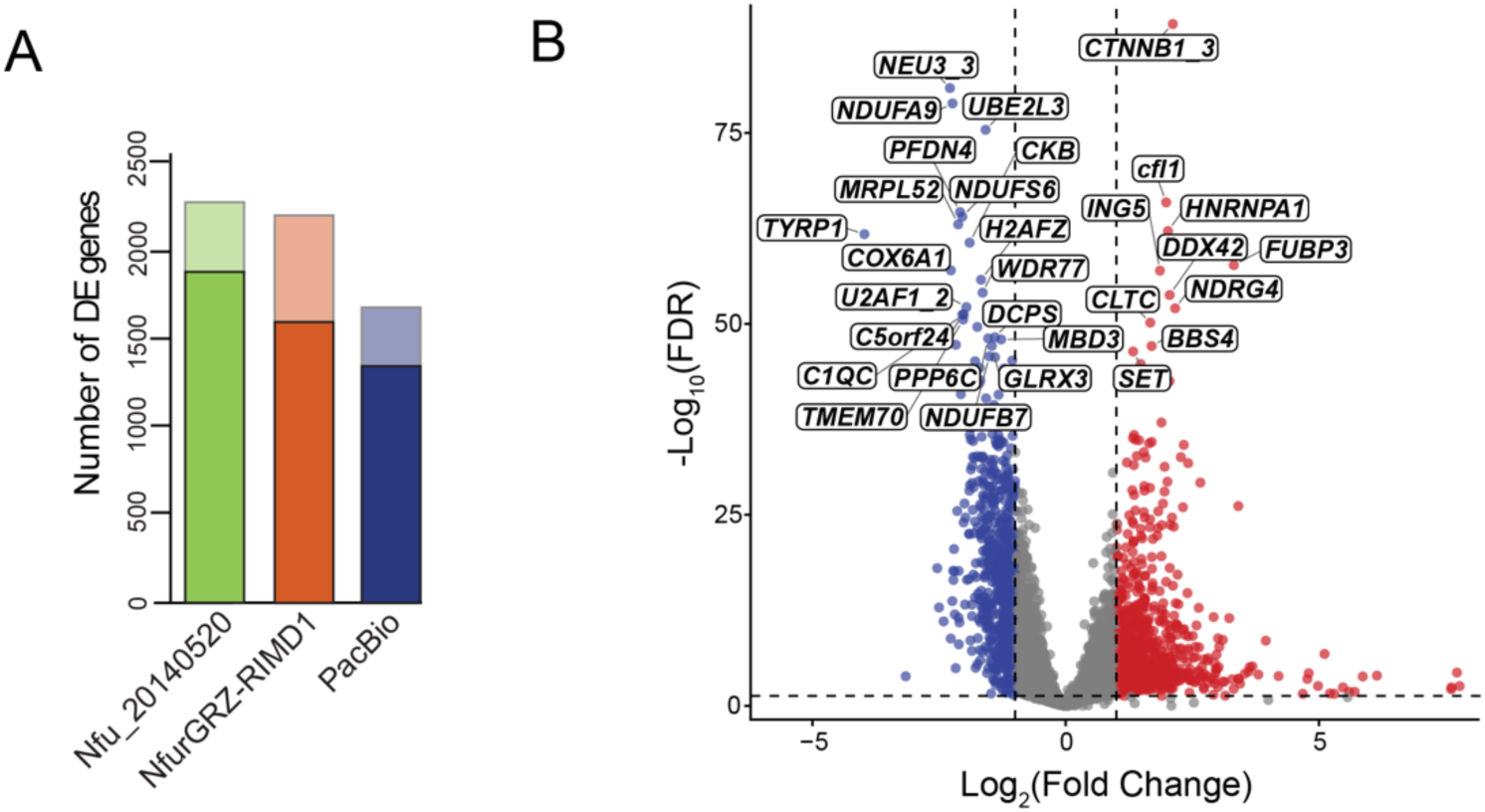
Improved Gene-Level Functional Annotation using the Pacbio Transcriptome. (**A**) Comparison of differentially expressed genes between 21- and 7-week-old retinal samples using Nfu_20140520, NfurGRZ-RIMD1 and PacBio transcriptome annotations. Bars represent the total number of DE genes and the proportion of biologically meaningful gene names (dark) and without gene name or placeholder gene names in (light). (**B**) Differential gene expression between 21- and 7-week- old retinal samples using the PacBio transcriptome, highlighting top 50 differentially expressed genes with biologically meaningful gene names (FDR < 0.05, |log2FC| >= 1).

**Table 1.** Comparison of gene and DE between 21- and 7-week-old bulk retinal across killifish transcriptome annotations.

| Transcriptome Build | Expressed genes retained, n (%) | Expressed genes with biological names (%) | DE genes | DE genes with biological names, n (%) |
| --- | --- | --- | --- | --- |
| Nfu_20140520 | 15,373 (57.5%) | 83.9 | 2,281 | 1,855 (82.6%) |
| NfurGRZ-RIMD1 | 14,757 (40.4%) | 73.2 | 2,207 | 1,600 (72.5%) |
| PacBio | 13,899 (66.1%) | 85.8 | 1,683 | 1,349 (80.2%) |

As the RNAseq dataset included multiple ages, we employed LOESS based clustering to identify global dynamics across time (**Figure 5A**, **Supplementary table 2**). Clusters 2, 3, and 8 were of particular interest as they showed consistent expression changes across time, which we hypothesize would be associated with age-related processes. GO analysis of the genes within each cluster identified shared and unique signatures (**Figure 5B**). Down-trending clusters 3 and 6 showing enrichment of mitochondrial and translational related pathways. Upregulated genes in cluster 2 were enriched for various cell signalling pathways such as GTPase regulators, transcription factors, and serine/threonine kinase activity. Identified genes and these cellular functions have previously been implicated in the aging process or linked to age related disease in other tissues and models [42–51].

**Figure 5:**
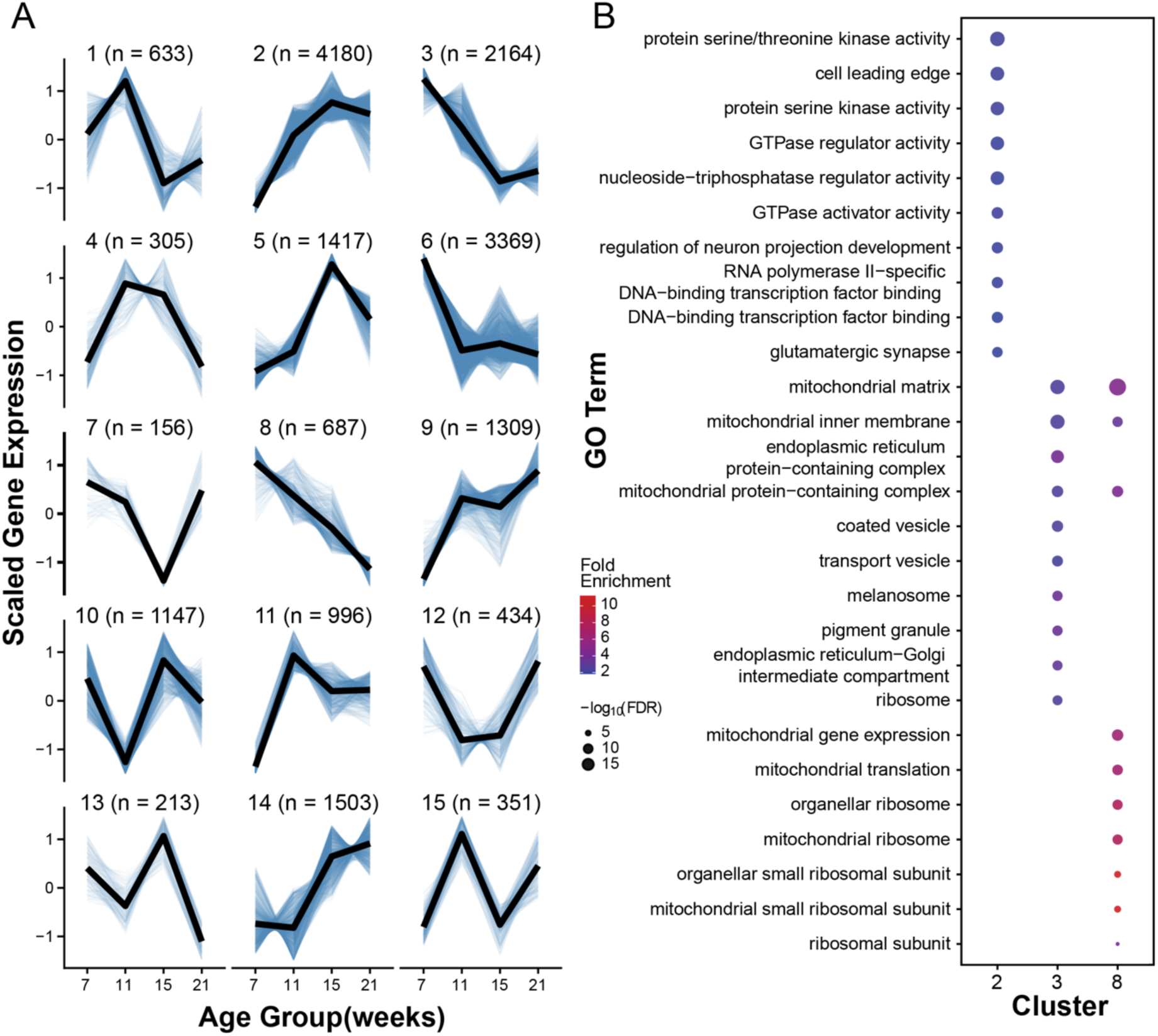
Age-Related Gene Expression Dynamics in the Killifish Retina. (**A**) Gene expression trajectories across retinal aging were identified using LOESS based clustering of scaled logCPM values. Thin blue lines represent individual gene trajectories, while the black line represents the average expression trajectory within each cluster. (**B**) Gene Ontology (GO) enrichment analysis of Clusters 2,3 and 8 associated with distinct age related expression patterns. Dot size represents the number of genes associated with each GO term, while color indicates enrichment significance.

Investigation of the DE list highlighted the potential loss of the green opsin encoding *rh2* gene (PB.6621) with age (**Figure 6A**). We noted that a second *rh2* gene (PB.6619) was located on the same chromosome 15.6kb away and showed high expression at all ages with no significant age-related dynamics. Indeed, a tandem duplication of opsin genes including *rh2* is common across teleosts [52–54]. It was notable to have both opsins properly named in our new transcriptome build as gene names in previous builds were unconventional or uninformative (Nfu_20140520: KFH-G, RH2-B; NfurGRZ-RIMD1: LOC107385170, LOC107385169).

**Figure 6:**
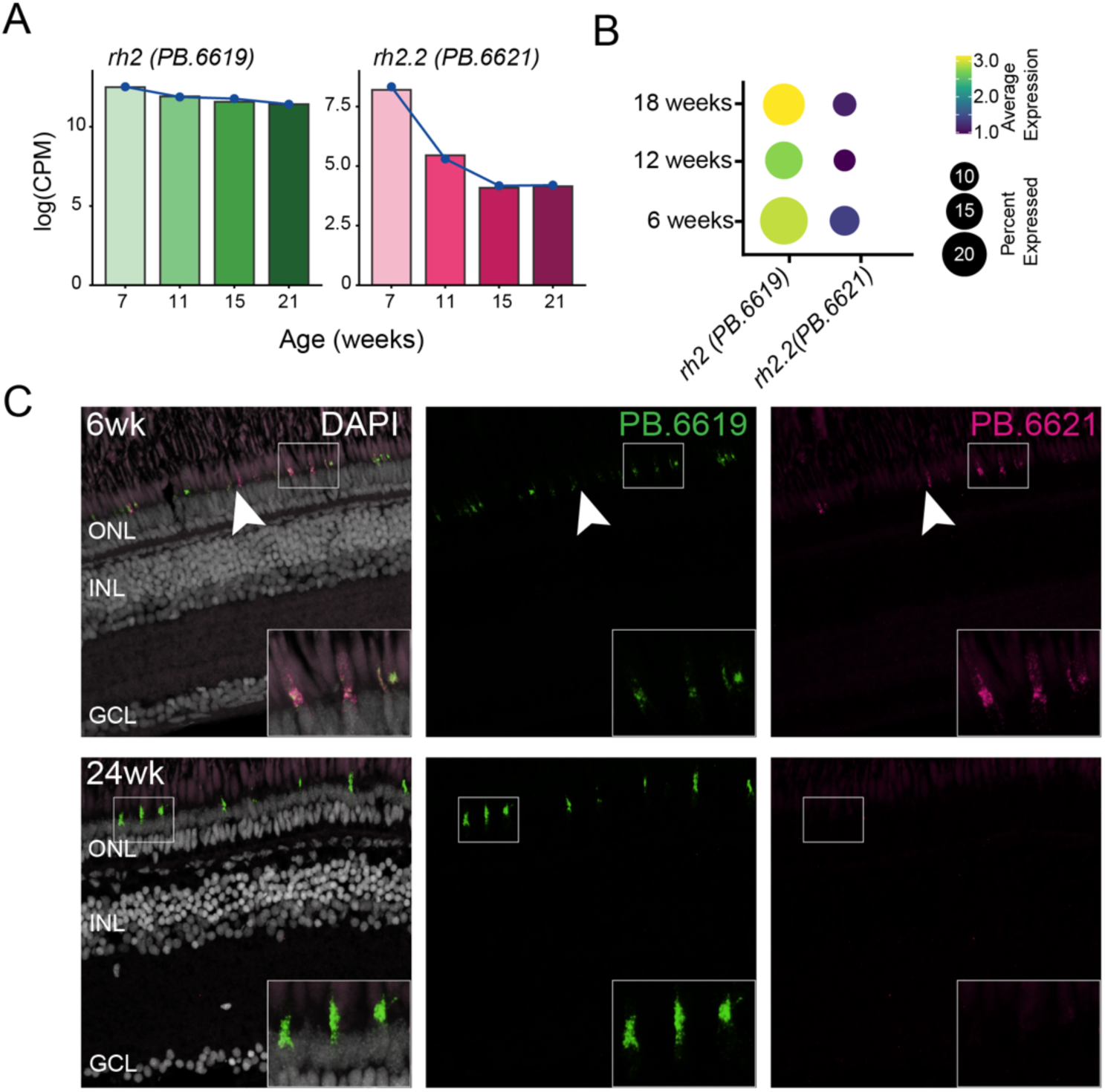
Age Associated Changes of Opsin expression. (**A**) Bulk RNA-seq expression of the two *rh2* genes, PB.6619 (green) and PB.6621 (magenta), across 7-,11-,15-, and 21-week-old retinal samples. (**B**) Dotplot showing expression of PB.6619 and PB.6621 in cone photoreceptors from remapped retinal scRNA-seq data at 6-,12-, and 18-weeks. (**C**) Hybridisation Chain Reaction (HCR) staining of 6- and 24-week-old killifish retina showing PB.6619 and PB.6621 expression. Arrow highlighting cells with independent expression of the two transcripts.

We first confirmed differential expression of *rh2* across time in the published scRNAseq dataset. Cone photoreceptors from 6-week-old killifish showed high expression of both *rh2* genes. Consistent with the bulk RNA expression data, *rh2* (PB.6621) expression decreased with time while PB.6619 maintained high expression (**Figure 6B**). Our scRNA data identified that the two *rh2* genes showed co-expression in the same clusters of cone photoreceptors (**Supplementary Figure 2B**). We sought to confirm both the co-expression and age-related loss of PB.6619 by Hybridization Chain Reaction (HCR) in killifish retina sections. Indeed HCR labeling of both *rh2* genes within young tissue showed co-expression in many cones (inset), though some independent expression was noted (**Figure 6C**, arrow). This pattern was lost in aged tissue where only signal from PB.6621 was observed, consistent with bulk RNAseq DE analysis.

## Discussion

A complete and well annotated transcriptome is an essential resource for any model organism. An incomplete or poorly annotated transcriptome can affect the analysis and limit biological interpretation of sequencing data and is a common issue for emerging model systems. These limitations are particularly relevant for the killifish which has become an increasingly useful vertebrate model for aging because of its short lifespan. However, transcriptome resources for killifish remain less developed, creating challenges for accurate identification of genes and transcripts that may change only subtly but have large impacts on the aging phenotype.

Our comparison of the available killifish genome and transcriptome resources highlights this problem. Although the newer NfurGRZ-RIMD1 genome displays improved genome completeness and assembly quality compared to the Nfu_20140520, its transcriptome annotation remains limited with nearly half of annotated transcripts lacking biologically meaningful gene names. We noted that both available reference transcriptome gtf files included a free-text attribute (‘product’ and ‘annotation’) that, in certain cases, contained a description of the orthologous protein even in the absence of a usable gene name. On an individual basis, a gene of interest could be hand annotated using these features. However, in practicality such descriptions are not beneficial for genome wide analyses.

By combining the improved NfurGRZ-RIMD1 genome with PacBio long read sequencing of retinal RNA we were able to expand transcript representation and also improve functional annotations. This new transcriptome reference provides a more complete and biologically interpretable reference for studying ageing and diseases of the killifish retina. Long read sequencing allowed us to identify retina specific transcripts/genes that were missing from existing annotations. Indeed, tissues of the central nervous system, including the retina, are reported to uniquely express isoforms not commonly seen in other tissues of the body [55–60]. Second, orthology based annotation increased the proportion of genes with biologically meaningful gene names. These steps dramatically improved downstream analysis using the PacBio transcriptome for both bulk and single-cell RNA sequencing. While our research is focused on retinal aging and disease, we did note that benefits of this resource extended to other neuronal tissues, though alignment rates across other tissues, including the endoderm-derived liver and gut tissues, were somewhat decreased. Additional effort will be required to comprehensively annotate the NfurGRZ-RIMD1 transcriptome for best utility across tissues, a task which has taken many years even for the human and mouse references.

A recent study similarly demonstrated the value of integrating long and short read sequencing data to improve identifying thousands of novel transcripts in telencephalon [25]. However, limitations in available killifish transcriptome resources remain, particularly for other tissues. Our study extends this approach to retina, using the newer chromosome level NfurGRZ-RIMD1 genome assembly as the genomic reference. In addition to increasing the retinal transcript representation, we used orthology based functional annotation to improve gene identification. Importantly, we also directly compared the Nfu_20140520, NfurGRZ-RIMD1 and the improved PacBio transcriptome annotations using both the bulk and single cell RNAseq datasets. This allowed us to evaluate not only whether the new transcriptome extended transcript and gene representation but whether those improvements translated into better performance and biological interpretation of existing retinal sequencing data. This analysis identified many genes that change expression in the retina with age. Genes we previously identified and validated - including *optn*, *nfkb2*, and *pcdh15 -* replicated in this new dataset, while we were also able to identify many other differentially expressed transcripts. Lastly, pathway analysis, which was restricted previously, was performed using the orthologous matches of significantly altered genes to identify additional gene sets and biological functions that may have important roles in vertebrate retinal aging.

Finally, we noted that our new transcriptome reference included previously unannotated important retina specific genes. We first confirmed correct expression patterns of these newly identified transcripts in scRNAseq data. We also noted that the PacBio transcriptome identified tandem duplication of the green-sensitive *rh2* opsin gene and bulk RNAseq suggested one of those genes loses expression in the aged retina. Analysis of the two *rh2* genes in scRNAseq data identified a cluster of cells that co-expressed the two opsin pigments. Duplication, co-expression, and changes over time of opsin encoding transcripts have been reported in other teleost species [52–54,61–63]. Using retinal tissue collected from young and old killifish, we confirmed both co-expression of the two genes as well as the loss of the more distal gene expression in aged retinas.

Together, this work has generated a new tool to enhance the study of aging and disease in the killifish retina. We have benchmarked this new transcriptome against other available resources and confirmed its application in multiple common analysis paradigms. Future efforts to generate a consensus reference with utility across all tissues is warranted to broaden the impacts of this important new model system.

## Supplemental Files

**Supplementary Methods:** Hybridization Chain Reaction probe sequences targeting *rh2* (PB.6619) and *rh2.2* (PB.6621).

**Supplementary Table 1:** Differentially expressed genes between 21-week-old and 7- week-old killifish retina (FDR < 0.05, |log2FC| >= 1).

**Supplementary Table 2:** LOESS based clustering of age-related gene expression trajectories across killifish lifespan.

## Acknowledgements

This work was supported by funding from the Carl Marshall and Mildred Almen Reeves Foundation (P.A.R. and B.S.C.), the National Institutes of Health (EY002687 to the Department of Ophthalmology and Visual Sciences), and Research to Prevent Blindness (Career Development Award to P.A.R. and unrestricted funds to the Department of Ophthalmology and Visual Sciences). We acknowledge contributions towards killifish maintenance costs from Moorfields Eye Charity Springboard Award (SB-24B-106) and BrightFocus Macular Degeneration Postdoctoral Fellowship (M2022002F) to Nicole Noel.

R.B.M was supported by a Biotechnology and Biological Sciences Research Council (BBSRC), David Phillips Fellowship (BB/S010386/1), and BBSRC grant (UKRI705).

A.M.K. was supported on a Moorfields Eye Charity PhD studentship (GR001503) to R.B.M. This manuscript is the result of funding in whole or in part by the National Institute of Health (NIH) Public Access Policy. Through acceptance of this federal funding, NIH has been given a right to make this manuscript publicly available in PubMed Central upon the Official Date of Publication, as defined by NIH.

---

**Supplementary Figure 1:**
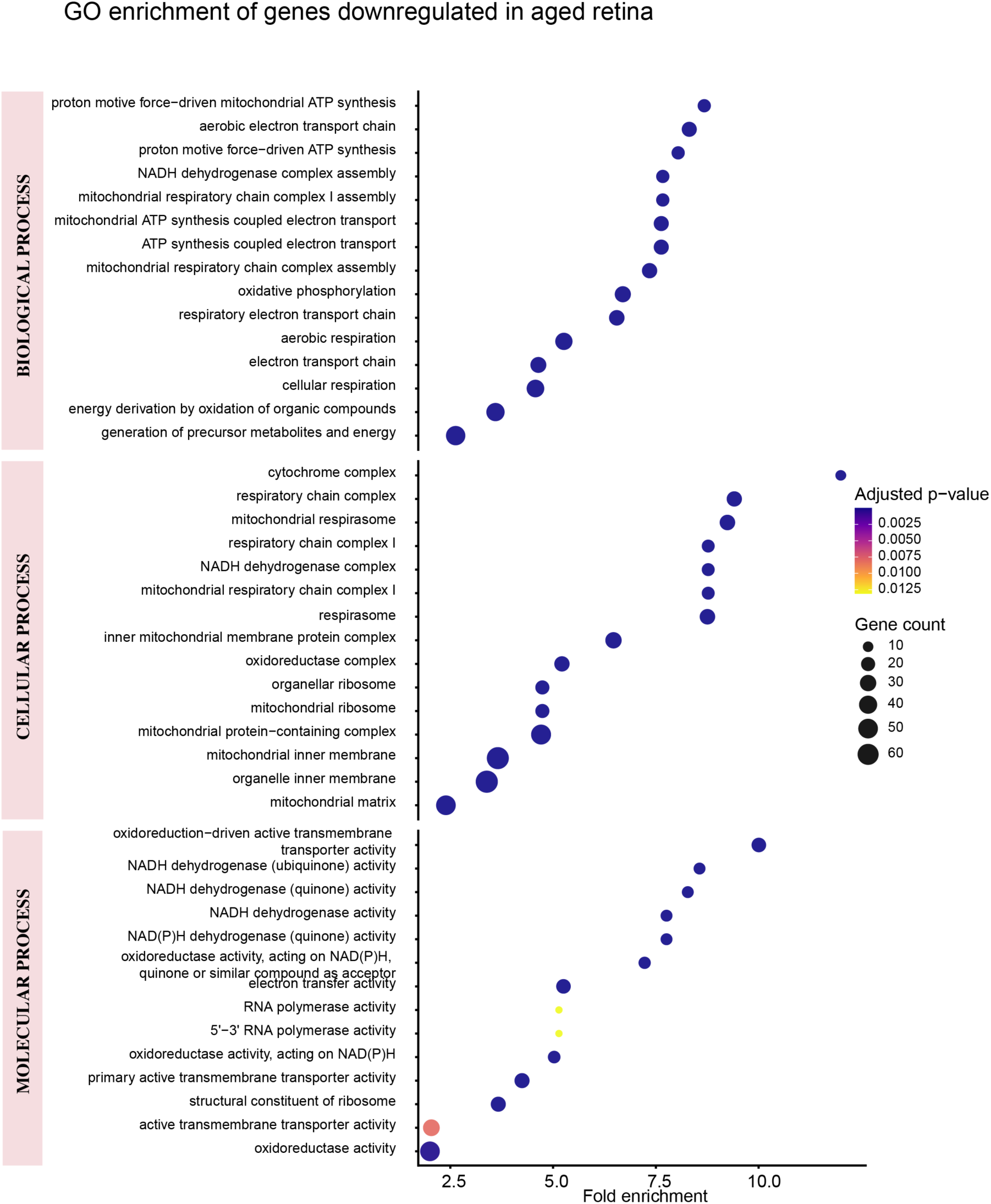
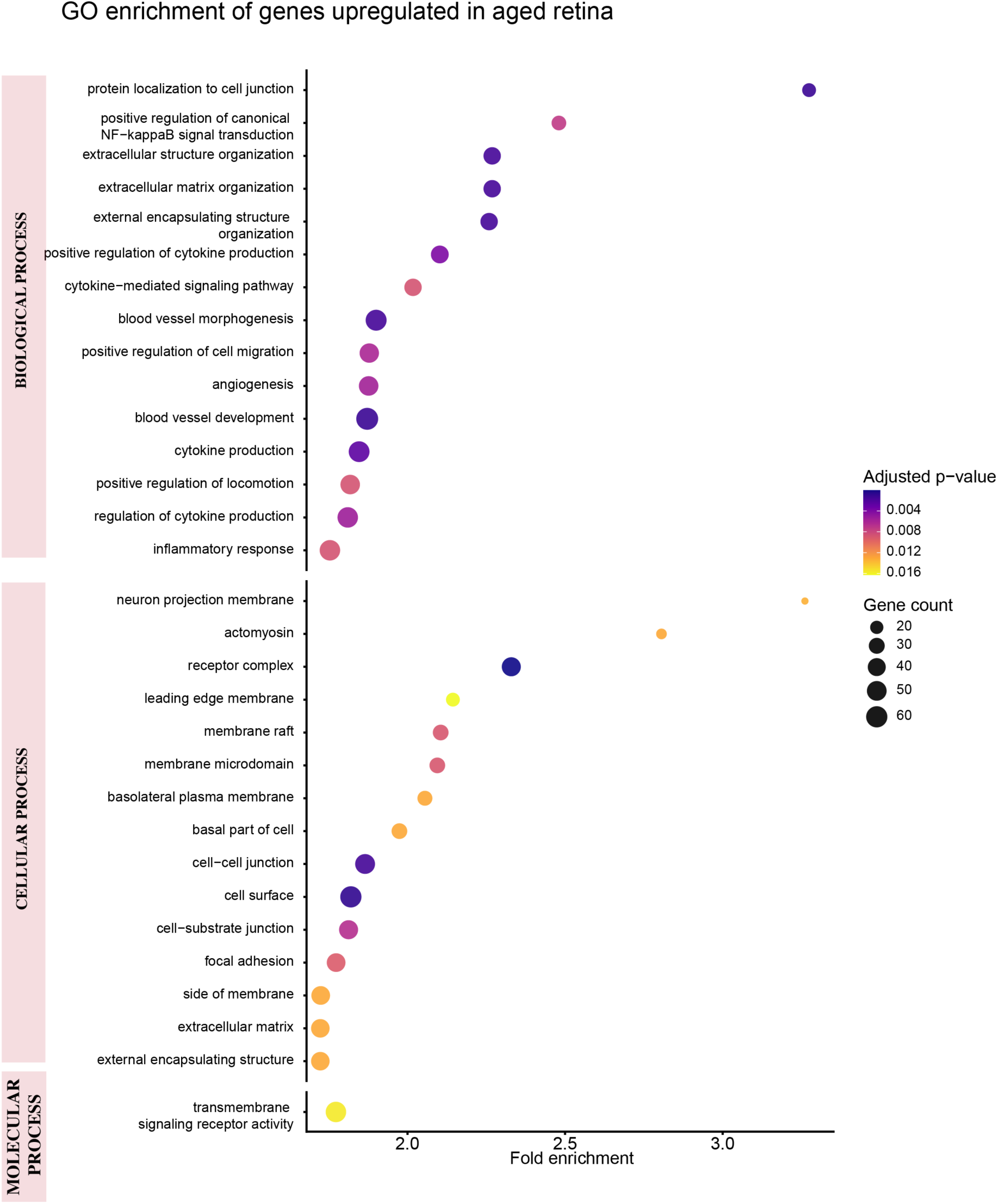
Gene Ontology analysis of age regulated differentially expressed genes in killifish retina (Top 15 enriched pathways). (A-B) Gene ontology enrichment analysis of genes significantly downregulated (A) and upregulated (B) in 21-week-old compared to 7-week-old retina.

**Supplementary Figure 2:**
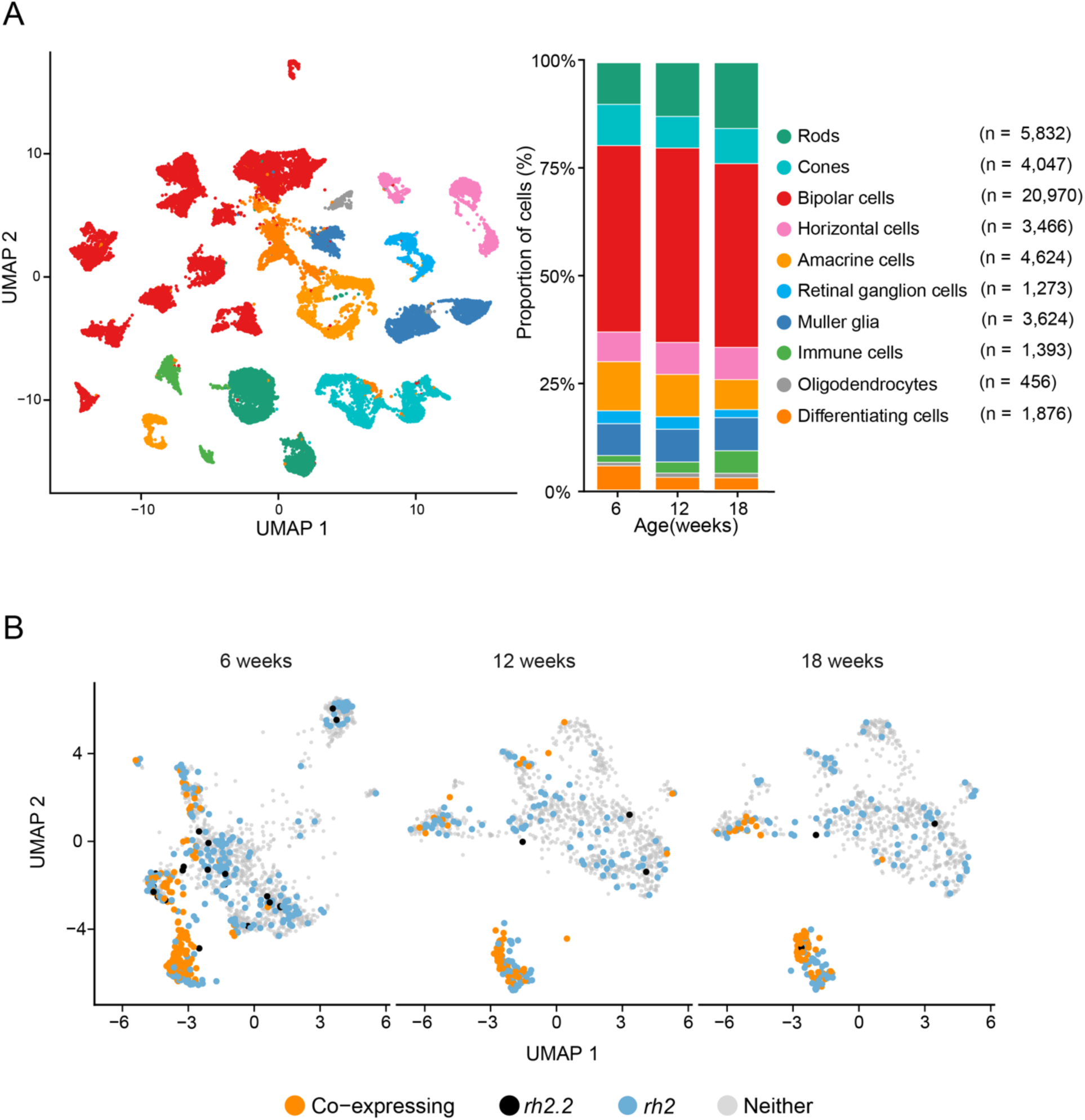
Cell recovery and *rh2* co-expression with PacBio transcriptome. (A) UMAP visualization and cell-type proportions across age groups. (B) UMAP visualization of *rh2* (PB.6619) and *rh2.2* (PB.6621) expression and co-expression in cone photoreceptors.

